# SSPIM: a beam shaping toolbox for structured selective plane illumination microscopy

**DOI:** 10.1101/253088

**Authors:** Mostafa Aakhte, Ehsan A. Akhlaghi, H.-Arno J. Müller

## Abstract

An important aim of the development of selective plane illumination microscopy (SPIM) is to present a completely open and flexible microscope set-up for nonspecialist users. Here, we report Structured SPIM (SSPIM), which provides an open-source, user-friendly and compact toolbox for beam shaping that can generate digital patterns for a wide range of illumination beams. SSPIM represents a toolbox to produce static, spherical Gaussian, Bessel and Airy beams by simple control of a Spatial Light Modulator (SLM). In addition, it is able to produce patterns for incoherent and coherent (lattice beam) array beam formation and tiling for all types of beams supported. We demonstrate the workflow and experimental and simulation results using the SSPIM toolbox. In final, the capability of the SSPIM is investigated with 3D imaging of *Drosophila* embryo using three different illumination beams such as scanned/dithered Gaussian, Bessel and Lattice beam which engineered with SSPIM. SSPIM toolbox is easy to use and applicable for a wide range of applications to generate and optimize the desired beam pattern and thus can help developing adaptation of the Open SPIM system towards a wider range of biological samples.

## Introduction

To understand the structure and function of living matter, scientists rely on imaging with conventional optical microscopy techniques. Optical microscopy easily permits the investigation of small details with a size of some micro meters to only a few hundred nano meters. While the natural environment for biological processes like the behaviour and interaction of cells in developing embryos happen within a three-dimensional space, a conventional optical microscope is only able to record a two-dimensional image of a sample. Therefore, a major goal of microscopy is to develop techniques that are able to obtain high-quality three-dimensional images of relatively large samples in their natural environment.

Selective plane illumination microscopy (SPIM) can be based on a simple open platform, where only a thin slice of a large object can be illuminated by a thin sheet of light [1] and the emitted light can be collected using a detection objective lens in perpendicular optical axes (Fig. 1a). SPIM was introduced with an static Gaussian beam (Fig. 1b) that is created with a cylindrical lens [1] which is called static light sheet [2]. Inside light scattering media, the beam will be diffracted and thus the propagation of the diffracted beam can produce some stripe artefact (ghost image) within the field of view (FOV) of the image. In order to reduce these artefacts, pivoting of the illumination beam was used [3]. Alternatively, a scanning circular Gaussian beam (Fig. 1d) can be used to overcome this problem [4]. One of the most important properties of the SPIM is its ability to image relatively large samples at a high resolution. For this reason, the illumination beam should ideally have a narrow and constant size during a large propagation length. The Bessel beam is one the optical beams, which has all of these parameters and, unlike the Gaussian beam, this beam consists of a narrow core surrounded with some rings (Fig. 1e). Fahrbach et. al. [2] used the scanning Bessel beam as illumination beam in light sheet microscopy and they proved that it can improve the FOV and reduce effects created by the ghost image. Following this work, other scientists have shown that the Bessel beam can be useful in single- or two-photon mode in SPIM [5]. However, the rings of the Bessel beam decrease the contrast and increase the out-of-focus effect. Gao et. al. have shown that by combining the Bessel beam with array beam which is called SIM mode, the contrast and axial resolution can be increased (Fig. 1h) [6]. In most cases, in SIM mode each plane of the sample should be recorded seven or nine times, hence, the imaging speed is slower than the conventional light-sheet mode. In order to increase the FOV with high axial resolution and imaging speed, the scanning and static Airy beam (Fig. 1c and 1f) was introduced which increase the FOV even more than a Bessel beam in SPIM [7]. Recently, a new generation of SPIM, called lattice light sheet microscopy was developed, which improved the axial resolution, reduced photo-bleaching and photo-toxicity when compared with other methods [8] (Fig. 1i). Most types of illumination beams that are used for SPIM can be conveniently generated by digital beam shaping methods using a spatial light modulator (SLM). In order to inform the beam shaping, the SLM requires some kind of digital pattern or digital mask.

**Figure 1.**
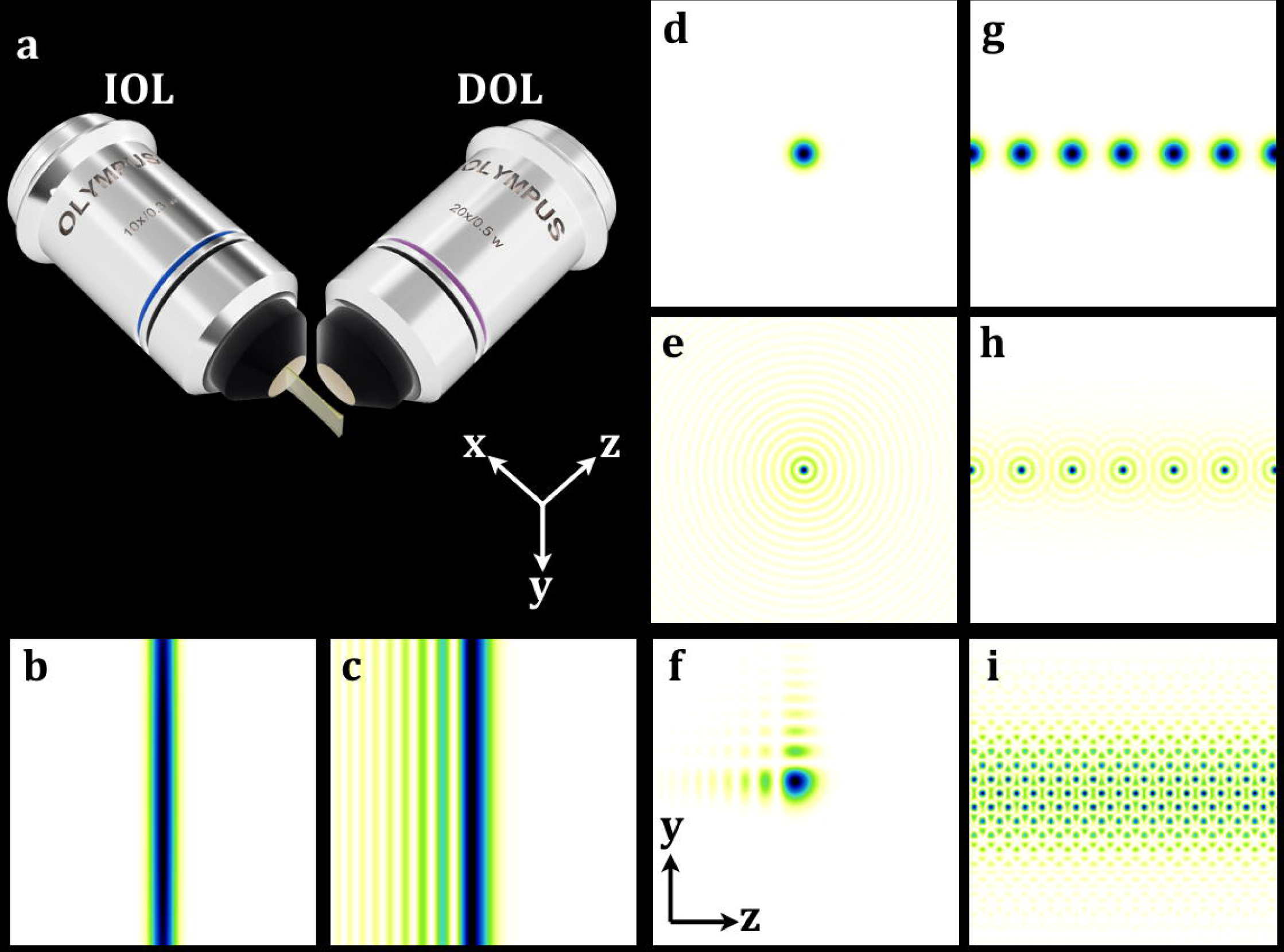
Schematic of the SPIM and illumination beams. Related to the size and transparency of the biological sample, one the illumination beam can be selected as an excitation beam. (a) Schematic of the conventional SPIM that the illumination objective lens (IOL) excites a thin sheet (x-y plane) of the sample, then the emission light is collected collects whit the detection objective lens (DOL) in the orthogonal direction (z axis). (b,c) the transverse intensity of static Gaussian beam and 1D Airy beam. (d,e) single Gaussian and Bessel with radial symmetry and (f) 2D Airy beams with asymmetrical intensity profile can be used for making a virtual light sheet. (g-i) in order to achieve better resolution and contrast, Gaussian and Bessel array beam and Lattice beams can be applied into the SPIM.

In this work, we present an open-source and user-friendly toolbox (SSPIM toolbox) for making digital phase mask or amplitude mask for beam shaping. SSPIM toolbox can support a wide range of the beams that have been routinely used in SPIM. SSPIM is designed with a graphical user interface (GUI) of MATLAB and is also able to work standalone. SSPIM has a user-friendly interface and easy to use for a biologist that has little knowledge in spatial laser beam shaping. Our goal is to help the SPIM community to improve towards more complete and flexible open SPIM systems.

## RESULTS

### Structure of the SSPIM toolbox

The toolbox is designed for generating a digital pattern for holographic beam shaping. The output of the toolbox can be used with a spatial light modulator (SLM) or it can be printed using other methods such as lithography. Output patterns of the SSPIM can be used with any type of SLM, such as gray scale or binary SLMs. It supports all types of beams that are currently used in light sheet microscopy. With the SSPIM, users are able to generate the digital pattern for static Gaussian beam, spherical Gaussian beam, Bessel Beam, 1D and 2D Airy beams. In addition, it can produce patterns for incoherent and coherent arrays of beams with three methods. For this purpose, Damman grating and optimal grating is used for incoherent array beam and a two-dimensional lattice method is used for generating a coherent array beam.

The work-flow of the SSPIM is composed of two major steps (Fig. 2). The first step is to define the wavelength of the illumination beam and spatial resolution of the SLM (Fig. 2a). In the second step, the operator has to select the type of the desired phase or amplitude mask. One of the following masks can be selected: (1) Cylidrical, (2) Spherical, (3) Annular, (4) 1D cubic, (5) 2D cubic and (6) lattice. With this selection, the phase or amplitude map of the desired beam will be calculated (Fig. 2b). For instance, if an annular mask is selected, a Bessel will be produced (Fig. 2c).

**Figure 2.**
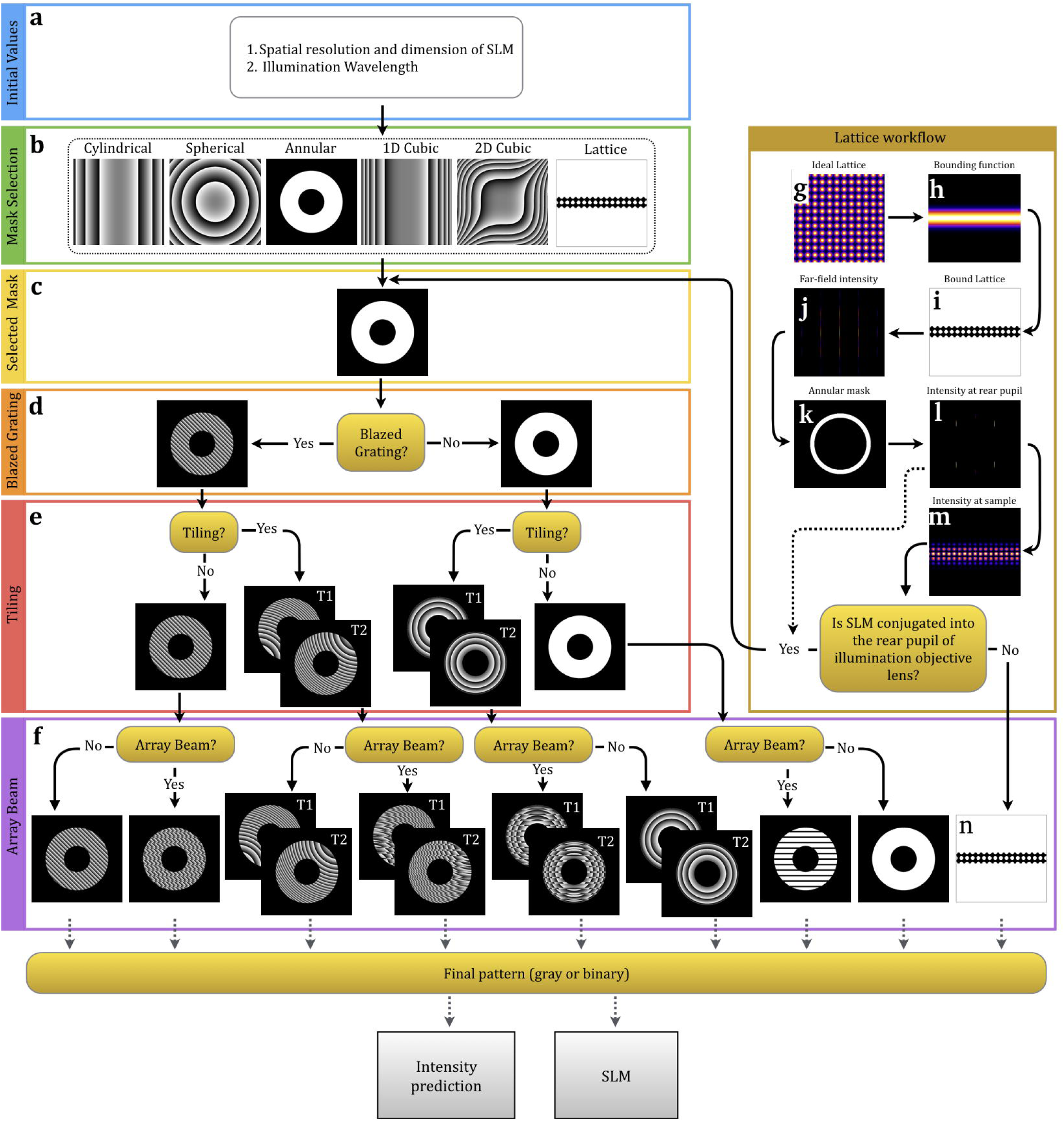
SSPIM workflow. (a) In the first step, toolbox is able to catch SLM parameters and illumination wavelength. (b) Then the desired beam can be selected from the six beam types. (c) As an example, the selected mask in this workflow is annular mask (Bessel beam). (d) If the optical setup needs to separate the diffraction orders of the SLM and hologram, a Blaze grating can be set (or binary grating for binary SLM). (e) Two, three or more quadratic phase can be added to the phase map in step (d), to tiling (swiping) the optical beam in the illumination direction. (f) In this step, user is able to add a binary Damman grating to the phase map to generating beam array. In step (b), if the user selects the “lattice”, another sub-menu will be opened for creating a SLM pattern for lattice beam. (g) An ideal 2D lattice (square and hexagonal), (h) bounding function and (i) bound lattice will be calculated. (j) Then the diffraction pattern of the bound lattice pattern will be predicted. (k) An annular mask can be set to select the desired spatial frequency of the bound lattice. (l, m) Then, the beam intensity at the rear pupil of IOL and in front focal plane of IOL will be predicted. If the SLM is conjugated into rear pupil, the bound binary lattice (i) will be selected as SLM pattern. Although, the calculated rear pupil intensity pattern (l) will be set as a selected mask. Then, all steps from (d) to (f) can be processed. In general, the output of the toolbox in step (f) can be used as an SLM pattern for beam shaping. The user is also able to truncate the phase map with an amplitude mask. In addition, the final SLM pattern can be saved for binary or gray-scale SLM.

An SLM has a pixelated structure meaning that the display of the SLM has active and passive areas, which enables it to generate a constant diffraction pattern like a two-dimensional grating. Hence, the diffracted light from the SLM is added to the diffracted phase or amplitude modulation and thus generates an unwanted diffraction pattern at the desired intensity. In order to separate these effects of the SLM, a Blaze or binary grating can be added directly to the phase or amplitude modulation, respectively [9]. The Blaze grating creates a linear phase with specific tilt angle and can change the centre of the desired beam on the Fourier plane. The transverse position of the desired beam can be adjusted in SSPIM by the orientation and spatial frequency of the Blaze grating (Fig. 2d).

One requirement to achieve high resolution and high contrast imaging using SPIM is the ability to generate a uniformly thin light sheet that is able to span the entire diameter of the specimen. In the case of the Gaussian beam, in order to increase the axial resolution, the waist of the Gaussian beam should be minimized, which diminishes the field of view (FOV). One of the best methods to enlarge the FOV with maximum axial resolution is tiling [10,11]. This method allows the illumination beam to swipe along the propagation direction. To achieve this, the selected phase map needs at least two quadratic phase maps for two tiling steps. In other words, the quadratic phase map is a spherical lens phase map, which the centre position of the beam can be determined with the focal distance (Fig. 2e). This tiling method can be used for single beam or array beam modes of the light sheet.

As mentioned above, a second mode of illumination beam in SPIM is a structured light sheet to recording high contrast and high resolution images. In principle, there are two ways to generate SIM light sheets. The first method is using a sinusoidal temporal light modulator and the second method is using a spatial light modulator (SLM) to creating array beam [6]. In order to create an array beam using a spatial modulation method we need an optical element that makes ‘n’ copies of the illumination beam. Damman and optimal gratings are diffractive optical elements, which split the incoming beam into a fan of ‘n’ outgoing beams with approximately similar intensity [13-16]. The number of the outgoing beams can be adjusted with transition number of the Damman grating structure and the angular spacing of the beams can be determined with the spatial frequency of the grating (Fig. 2f). The Damman or optimal grating can be merged with the hologram that was created in the previous steps for a single beam. It should be noted that these types of gratings generate an incoherent beam array meaning that the beams do not have specific phase relationship. In order to avoid any interference pattern between the beams, the angular spacing between the beams should be sufficiently large to avoid extensive overlap between the individual beams.

In this way to achieve an array beam with minimized undesirable out of focus light, lattice beam offers a high confinement array beam around the focal plane. In other hand, lattice light sheet microscopy is the most powerful method to perform high-resolution imaging with minimal photo-bleaching over a large field of view. The lattice beam is a structured beam, which is generated by superimposing coherent wave planes in well-defined directions and can be used in two modes of light sheet illumination: dithering (scanning) and SIM mode [8].

The lattice beam is a multi-core non-diffracting beam like a Bessel beam. The intensity profile of the beam is invariant over the specific propagation length. For generating a lattice beam with maximum light confinement, the SLM pattern should be carefully engineered. For this purpose, first, the desired 2D lattice should be defined. In this step, the structure of the lattice, size of the spots and distances to each other can be defined. SSPIM toolbox is able to generate two types of lattice beams, either square or hexagonal (Fig. 2g). In second step, for generating a 1D-like lattice beam a bounding function with an arbitrary width can be applied (Fig. 2h). In third step, for generating coherent Bessel beam array, the diffraction pattern of the 1D lattice (Fig. 2j) can be adjusted with an annular mask in Fourier domain (Fig. 2k-l). The intensity prediction at this step (Fig. 2l) shows the intensity at rear pupil of the objective lens. Finally, the intensity profile of the desired lattice beam at sample place will be predicted and shown with SSPIM (Fig. 2m).

It should be mentioned that if the SLM is conjugated to the front focal plane of the illumination objective (Supplementary Fig. S1b), the SLM pattern can be extracted directly from the bound lattice (Fig. 2i or 2n). Although, if the SLM is conjugated to rear pupil of the illumination objective lens (Supplementary Fig. S1a), the SLM pattern is equal to the binarized Fourier transform of the bound lattice (Fig. 2l).

### Laser beam shaping using SSPIM

Our toolbox is designed for two types of SLM, the binary and the gray scale SLM. A fast binary SLM is implemented in our light sheet SPIM for high-speed imaging. The binary SLM is conjugated into the back focal plane of the objective illumination lens (Supplementary Fig. S1a). Hence, the Fourier transform of the hologram can be achieved in the front focal plane of the objective illumination lens. The SSPIM toolbox can be used for static and for scanning beams. We examined the ability of the SSPIM to generate desired patterns for beam shaping by imaging the beam intensity in cross-section and over the propagation. In order to generating static Gaussian beam and 1D Airy beam, quadratic and cubic phase modulation can be used, respectively (Fig. 3a1 and 3b1). The beam waist and the Rayleigh lengths of the static Gaussian beam can be adjusted with the rectangular amplitude mask. As well, the invariant propagation length and stretching of 1D Airy beam in y direction can be controlled with cubic phase coefficient and the size of rectangular mask, respectively (see **Methods**).

**Figure 1.**
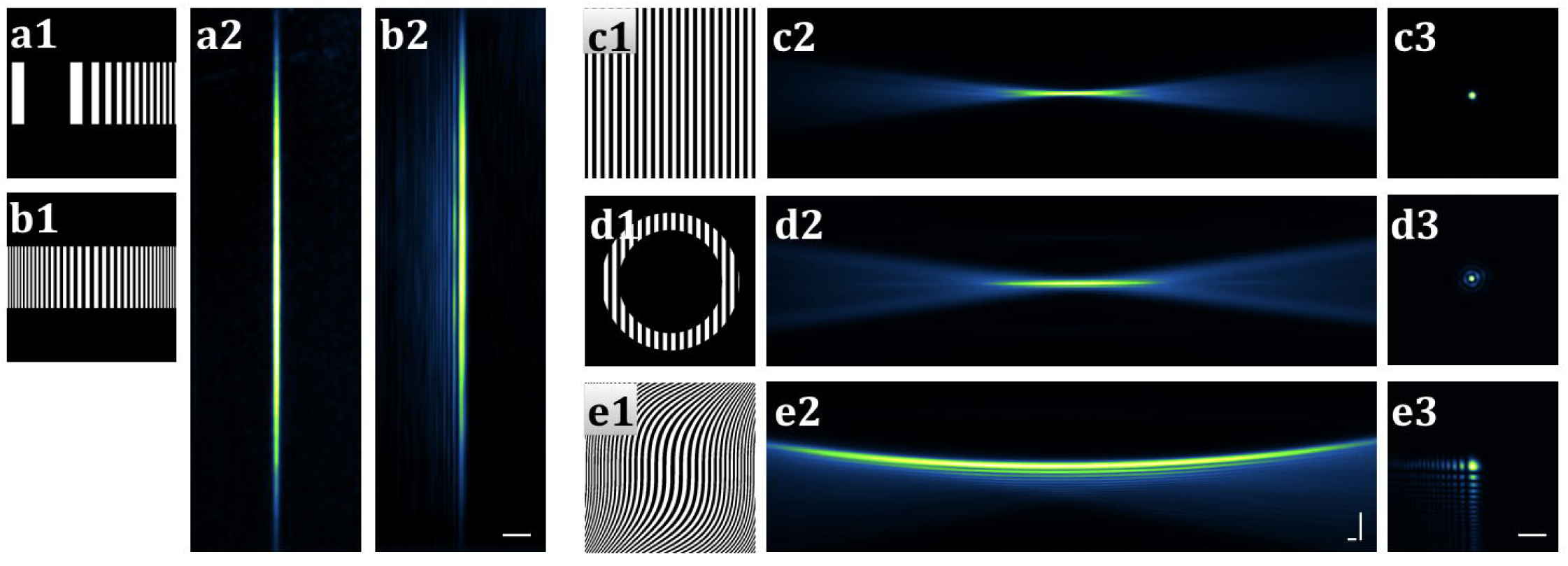
SSPIM output for single beams. (a1-a2, b1-b2) The binary pattern and recorded cross-section intensity of the static Gaussian and 1D Airy beam, respectively. (c1, d1 and e1) The calculated SLM patterns to create the circular Gaussian, Bessel and 2D Airy beams. (c2, d2 and e2) Measurement of the beam propagation intensity through the dye solution. (c3, d3 and e3) Recorded the cross–section of the beams. Scale bars: 20 μm.

The output of the laser that we used in these experiments has a Gaussian beam with circular symmetry. Therefore, to using Gaussian beam as an illumination beam, we need use only a binary grating for separation of the Gaussian beam from the default SLM diffraction pattern (or blazed grating for gray value SLM) (Fig. 3 c1-c3). A wide range of the Rayleigh lengths of the Gaussian beam can be produced with changing the radius of the circular amplitude mask (Supplementary Fig. S2). On the other hand, by decreasing the radius of the amplitude mask (Supplementary Fig. S2. a1-d1) the numerical aperture and the Rayleigh length of the Gaussian beam is decreased and increased, respectively (Supplementary Fig. S2 a2-d2).

For high-resolution imaging of a large specimen, the SPIM demands illumination beam that offers a thin main lobe with a large invariant propagation length like the Bessel beam. One method to generate a Bessel is using an annular mask (Fig. 3d1-d3). The properties of the Bessel beam can be adjusted by changing the inner and outer radius of the annular mask (see **Methods**). The core size of the beam and the number of the surroundings rings can determine the axial resolution, contrast and the field of view of the image. One method to achieve a smaller core of the Bessel beam with increasing the number of rings, is to proportionally increase the inner and outer radius of the mask (Supplementary Fig. S3 a1-c4). Alternatively, the Bessel beam can be manipulated with a constant number of rings to minimize the propagation length and size of core (Supplementary Fig. S3 d1-f4). These capabilities are critical to adjusting the parameters of the Bessel beam with the dimensions of distinct specimens.

Another type of beam that was more recently introduced in SPIM applications is the Airy beam. Airy beam is another type of non-diffracting beam that is able to conserve the intensity along the specific propagation length. Although illumination with the Airy beam somewhat reduces the axial resolution, it was introduced because it significantly improves the FOV of the imaging. It is well known that different cubic coefficients can generate different 2D-Airy beams in which the trade-off between an increased propagation and a decreased axial resolution has been observed (see **Methods**). In order to generate a 2D-Airy beam SPPIM toolbox applies a two-dimensional cubic phase (Fig. 3, e1-e3). SSPIM supports the application of different cubic coefficients to modify the size of the main lobe of the Airy beam and thus enables the user to optimize for the properties of the specific specimen with regard to axial resolution and FOV (Supplementary Fig. S4)( Supplementary Movie1).

SSPIM also has the capability to produce both coherent and incoherent beam arrays. Multiple incoherent copies of the incident beams can be generated by a binary Damman grating also known as a ‘fan-out element’. SSPIM can generate distinct Damman grating holograms supporting the formation of array beams composed of 2 to 21 individual beams. For example, a 7 Gaussian beam array can be produced with its characteristic propagation and cross-section intensities (Fig. 4 a1-a3). The Bessel beam array also can be achieved easily by merging an annular mask with Damman grating (Fig. 4 b1-b3). The propagation and cross sections of Gaussian and Bessel array beams with different array number were recorded in dye solution and compared to results from simulations (Supplementary Fig. S5 and S6). SSPIM contains an array beam sub-menu which the angular spacing and number of the spots can be easily selected (Supplementary Movie2, Movie3).

**Figure 2.**
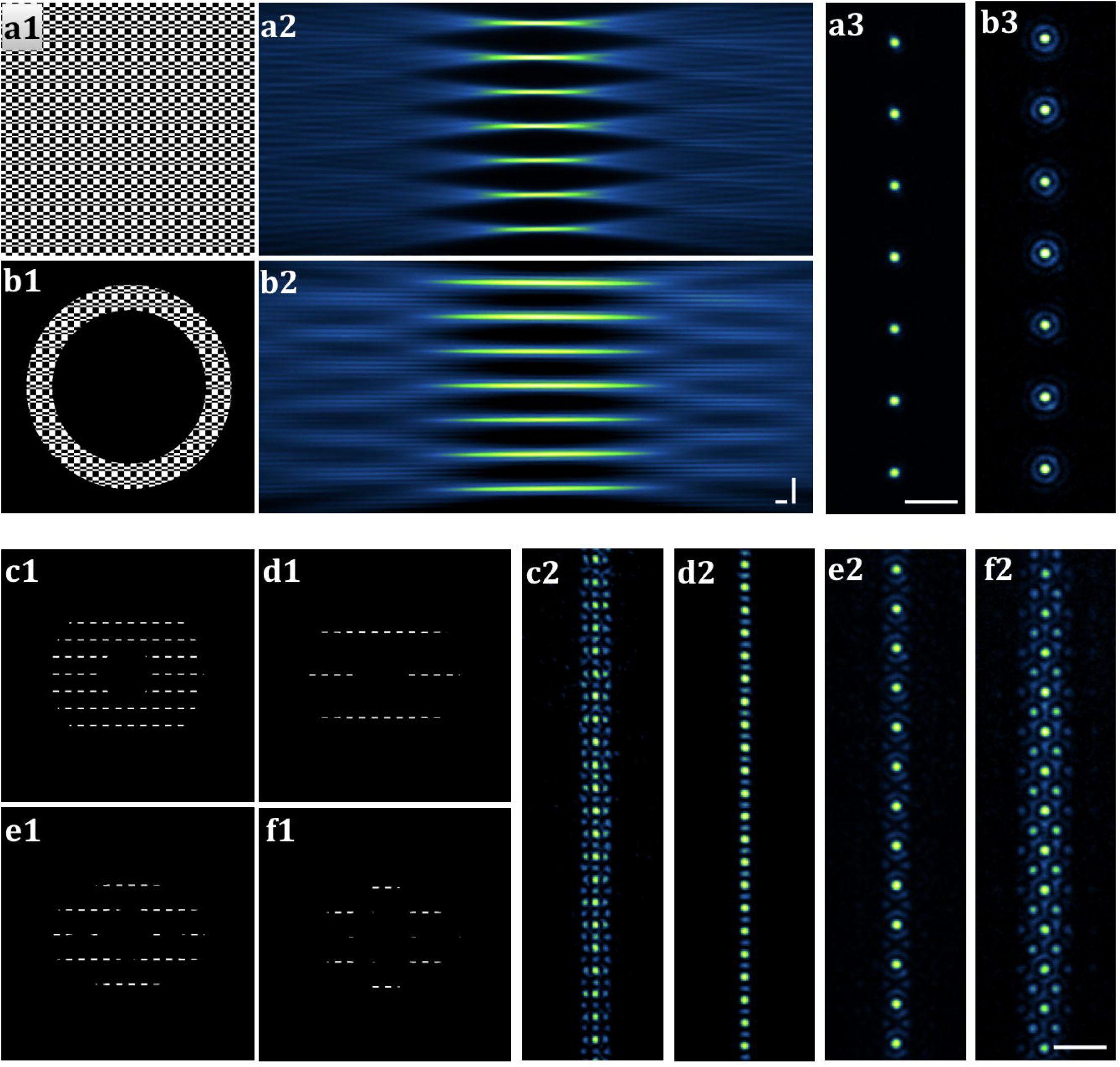
SSPIM output for Array and Lattice beams. (a1, b1) The binary pattern applied to the SLM to generate the 7-array of Gaussian and Bessel beams. (a2, b2) Recorded propagation intensity of the Gaussian and Bessel beam array through the dye solution. (a3, b3) Measurment of the cross-section array beam intensity. (c1, d1) Created SLM patterns to generate square lattice beams with different bounding function. (c2, d2) Measured cross-section intensity of square lattice beams. (e1-f1) Created SLM pattern to generate hexagonal lattice beam with different bounding function. (e2, f2) Measured cross-section intensity of hexagonal lattice beams. Scale bars: 20 μm.

As mentioned above, the SSPIM toolbox also supports an SLM pattern for coherent lattice beams. In this case, the toolbox has two options for users that is related to the position of the SLM in setup (Supplementary Fig. S1a and S1b). In our case, the SLM was conjugated into the rear pupil of the objective lens. Hence, the Fourier transform of the SLM pattern can be obtained in front focal plane of the illumination objective lens. The Lattice sub-toolbox is designed to produce square and hexagonal patterns. To evaluate the ability of this feature in SSPIM, we generated four different SLM patterns and analysed the cross-section intensity of the beam (Fig. 4c1 – 4f2). The SSPIM toolbox can readily produce different types of lattice SLM patterns, which can then be used in the conventional arrangement of an Open SPIM system (Supplementary Fig. S1). The properties of the lattice beams such as angular spacing of the intensity spots, size of the spots, and beam confinement can be adjusted easily with SSPIM (Supplementary Fig. S7 and Fig. S8) (Supplementary Movie3, Movie4 and Movie5).

### SSPIM application for tiling in deep tissue imaging

The live imaging of dynamic cell and tissue movements in relatively large sample requires a very thin optical beam with large FOV. Tiling is a relatively simple method to approach this challenge. With the tiling method, the excitation beam can be swiped along the illumination direction, and therefore can generate a thin light sheet with a large FOV. The SSPIM toolbox allows the user to apply tiling on the conventional SPIM and it supports the generation of tiled holograms for all types of beams that we have introduced. We created two types of the three tiled holograms, for the single and array of Gaussian beam (Fig. 5a and 5b). The results demonstrate that coverage of a specific FOV is easily accessible with the tiling of an optical beam with small invariant propagation length. The precision of axial resolution and enlargement of FOV can be controlled with the numerical aperture of the illumination beam and number of tiles (Fig. 5 (a1-a6)). The combination of Damman grating with tiled phase maps can produce tiled array beams which can be applied in light sheet-SIM (Fig. 5 (b1-b6)). To examine the capability of SSPIM to support the tiling method for all types of beams, we created tiled holograms and recorded the intensities of the tiled beams (Supplementary Fig. S9). The results show the tiling method can be applied successfully for all beams like as array Gaussian, array Bessel and lattice beam. All of the pattern that we used for tiling are binary and it should be noted that the SSPIM is able to support the tiled gray pattern for gray SLMs.

**Figure 3.**
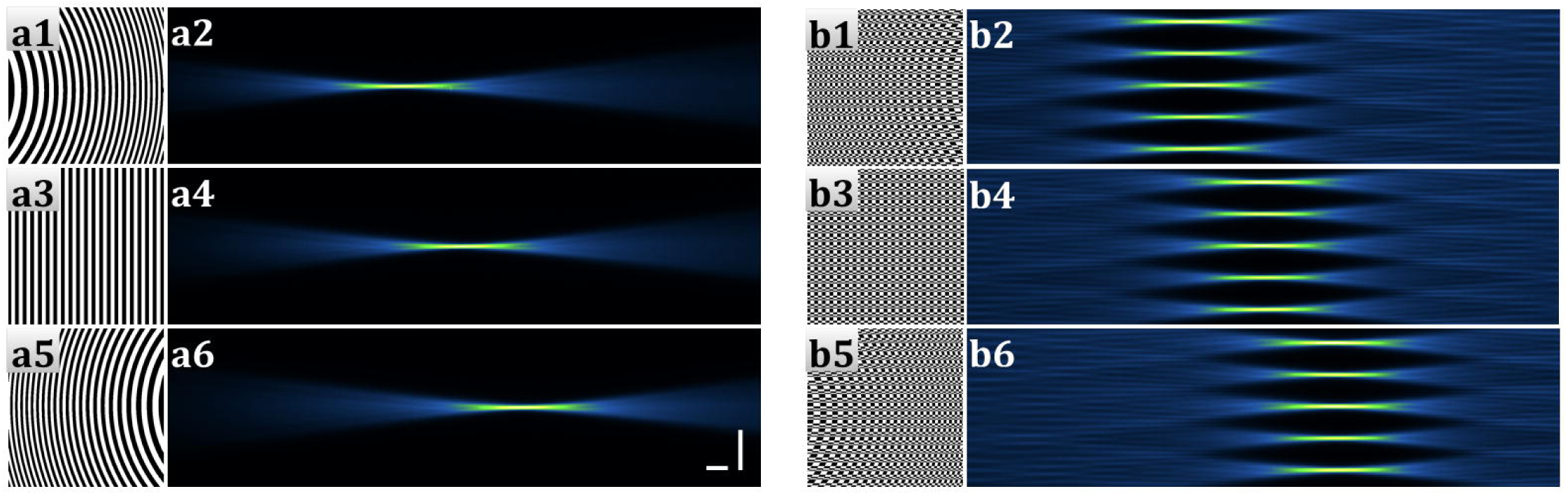
SSPIM output for Tiling method. (a1, a3, a5) SLM patterns to tiling a single beam in three steps. (a2, a4, a6) Recorded the propagation of the tiled beams. (b1, b3, b5) SLM patterns to tiling a 5 beam array beam in three steps. (b2, b4, b6) Recorded the intensity propagation of the tiled beam array. Scale bars: 40 μm.

### SSPIM application in three-dimensional imaging with light sheet-SPIM

Engineering of the illumination beam is an essential way for improvement of the axial resolution, field of view and the time resolution in light sheet-SPIM. In order to examine the capability of SSPIM toolbox we recorded a set of images of a fixed sample (*Drosophila* embryo) using Gaussian, Bessel and square lattice sheet (Fig. 6). The SLM patterns were engineered with SSPIM such that the same FOV was recorded. Recorded images of the specimen in the axial direction are analyzed in Fourier domain. Obviously, in frequency domain the cut-off radius in axial direction of the data using lattice sheet is approximately 40% and 30% larger than the Bessel and Gaussian sheet, respectively (Fig. 6b). It means, the spatial resolution of recorded image in axial direction using lattice beam is better than the two other beams. The comparison of the deconvolved data in spatial domain using these sheets show the recorded image using lattice sheet can resolve more details of the sample in axial direction (Fig. 6c). The properties of these beams such as invariant propagation length, beam waist, and the number of the array beam were readily adjusted with SSPIM.

**Figure 4.**
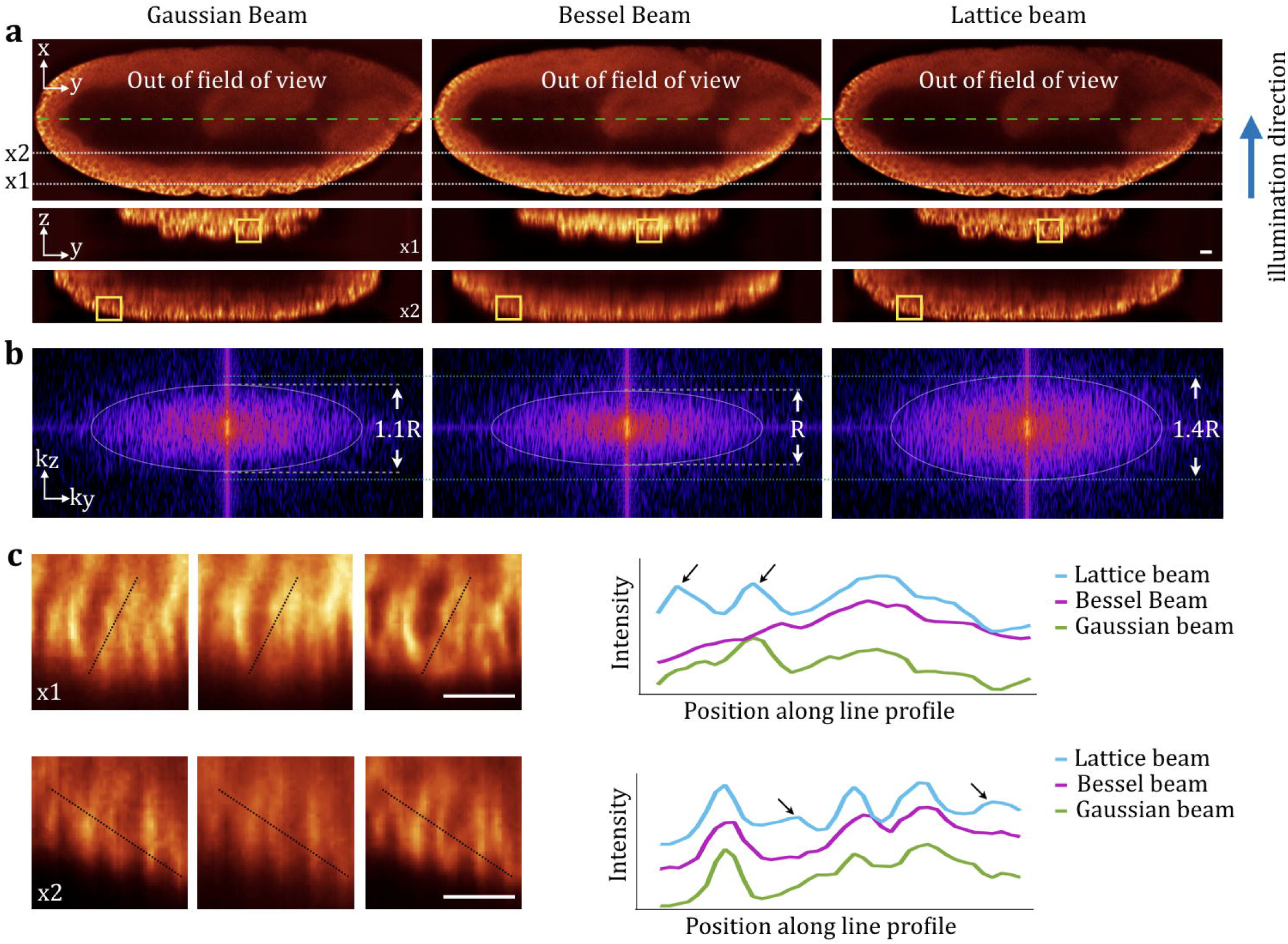
Resolution improvement with engineered illumination beam using SSPIM. (a (first row)) Recorded images of Drosophila embryo with SPIM using scanned Gaussian, Bessel beam and dithered square lattice beam. (a (second row)) Axial view of recorded data in different depth (x1 and x2). (b) Improvement of spatial resolution can be achieved with using engineered beam like as lattice beam. Fourier analysis of the axial view of recorded images show the radius of cut-off frequency is increased approximately 40% with lattice beam than to the Gaussian and Bessel beam. (c) Enlarged view and line profile of axial views in (a) show increase the resolution using lattice beam. Black arrows show the difference area that resolved by lattice beam that were not resolved with Gaussian and Bessel beam. Scale bars: 15 μm.

## Discussion

In this work, we introduced SSPIM toolbox which can be used for holographic beam shaping for light sheet - selective plane illumination microscopy. The toolbox presented here is open-source and user-friendly for non-specialist users. It covers most of the optical beams for wide range applications. With SSPIM, a user is able to design a digital pattern for a specific resolution, field of view and light confinement. The desired pattern can be easily generated for binary and gray value SLM and the toolbox is able to simulate the intensity of the designed pattern (hologram). When using SSPIM it is important that the illumination beam should be carefully selected to achieve a desired spatial resolution, light confinement and field of view of a specific sample. Some of the important biological specimens that imaged with SPIM or Light sheet are summarised in table 1. To examine the capability of the SSPIM in simple SPIM setup we have created three different SLM pattern for generating Gaussian, Bessel and lattice sheet. These beams are successfully tested for imaging using a fixed biological sample (*Drosophila* embryo). The results show that the resolution was improved by engineering the optical beams with SSPIM toolbox. As well the SSPIM toolbox has capability to work in lattice light sheet mode in conventional SPIM without need to change structure of the microscopy setup. The SSPIM toolbox will allow the user to examine different beam profiles on distinct biological samples to optimise imaging of a specific specimen using the open SPIM platform.

**Table 1.** List of biological samples which have imaged with different type of beams.

| Illumination beam/Illumination method | Biological samples |
| --- | --- |
| Elliptical beam (static) | *Medaka* embryo [1], *Drosophila* embryo [1], Zebrafish embryo [20]. |
| Gaussian Beam (scanning mode) | Zebrafish embryo [4], *Drosophila* embryo [21]. |
| Bessel Beam (scanning mode) | mEmerald, U2OS cell, HeLa cell [5], |
| 1D Airy Beam (static mode) | zebrafish larva[22], BY-2 cells, rat brain [23]. |
| 2D Airy beam (scanning mode) | Juvenile amphioxus, renal adenocarcinoma cell ,zebrafish [7], non-cleared and cleared mouse brain[24]. |
| Gaussian beam array (SIM mode) | *Drosophila* embryo [12]. |
| Bessel beam array (SIM mode) | LLC-PK1 cell , *Dictyostelium discoideum* cell and *C. elegans* embryo[6]. |
| Lattice beam array (Dithering and SIM mode) | HeLa cell, *D. discoideum* cell, mouse embryonic stem cell, U2OS cell, mEmerald-Lifeact, *C. elegans* embryos, *Drosophila* embryo[8]. |
| Tiling method (Gaussian beam, Bessel beam, Lattice beam) | *C. elegans* embryo, zebrafish embryo[10,11]. |

## Methods

### SSPIM availability

The SSPIM toolbox and the associated documentation are available at https://github.com/aakhtemostafa/SSPIM. The toolbox runs under the popular MATLAB computational environment and standalone on Windows and Mac.

### Open SPIM setup

A single illumination-detection light sheet-SPIM was built for the research (Supplementary Fig. S1a). A CW laser source (Coherent OBIS 488 nm) was used for linear excitation. The laser beam was expanded to 8.5 mm diameter (L1=25 mm, L2=300 mm) and sent to a fast binary spatial light modulator (SXGA-3DM Forth Dimension Displays) for beam shaping. The binary SLM assembly, consisted of a half-wave plate and a 1280×1024 binary SLM, was used to modulate the wavefront and amplitude of the excitation light to generate different light sheets such as Gaussian, Airy, Bessel, coherent array and incoherent lattice beam. An Iris was placed at the focal plane of the lens L3 after the SLM to block the undesired diffraction orders of the excitation light generated by the SLM (L3=150 mm, L4=75 mm). After the SLM, the laser beam was sent to the beam scanning or dithering mirror to create the light sheet (10 mm Galvanometer mirrors, Thorlabs, GVS212/M). The Galvo mirror and SLM were conjugated to the rear pupil of the illumination objective lens (Olympus UMPLFLN 10XW 0.3 NA). In order to collect 3D data, the sample was moved through the light sheet with a piezo (Q-522.040, PI), While it was embedded into 1.5% agarose gel that held with a glass capillary. The emitted fluorescence was collected with a detection objective lens (Olympus UMPLFLN 20XW 0.5 NA), was placed orthogonal to the illumination objective lens and was imaged onto a sCMOS detection camera (Hamamatsu, ORCA-Flash 4.0 V2) with a tube lens (U-TLU-1-2 tube) and a fluorescent filter. The SPIM setup was controlled with a home-built software that is built by Labview, and the final data were deconvolved and reconstructed with Lucy-Richardson method using MATLAB and Fiji.

### Cylindrical Lens phase map (Static Gaussian beam)

A static light sheet can be produced with a cylindrical lens and the transmittance function of a cylindrical thin lens can be defined:

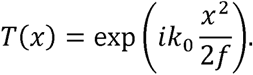

Where the *k*_0_, *x*, and *f* show the wave-number, transverse coordinate, and focal length [17]. If assume the incident beam has plane wave, intensity distribution in the focal plane of the cylindrical lens looks like a line. Hence, propagation of the line beam can make a static sheet of light in Rayleigh range.

### Spherical Lens phase map (circular Gaussian beam)

In order to achieve a focused beam with Gaussian intensity distribution, we can use a spherical lens. In contrast to cylindrical lens, Fourier transform of the incident -beam in the radial direction can be achieved with a spherical lens. Since the intensity distribution in the focal plane is a Gaussian spot with radially symmetric properties. The transmittance function of a spherical lens can be written as

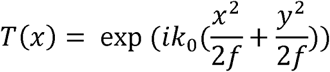

where, *x* and *y* denotes the transverse coordinates, *k*_0_ is the wavenumber and *f* shows the focal length of the lens [17].

### Bessel and Airy Beams

Another type of the optical beams that were introduced for selective plane illumination microscopy are non-diffracting beams such as Bessel and Airy beams. The Bessel beam can be generated with conical superposition of the plane waves. The angular spectrum of a Bessel beam can be approximated by a ring. Because, the Fourier transform of a ring with inner radius *R_i_* and outer radius *R*_0_ is zero order of a Bessel function. Thus, the Bessel beam can be created at the front focal plane of a Fourier lens, where an annular mask is placed in the back focal plane of it. The annular aperture can be defined with this amplitude mask:

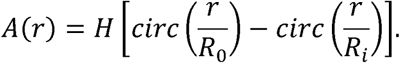

The *H* denotes the Heaviside function, *circ* is the circular function, *r* is radial transverse coordinate, *R*_0_ and *R*_i_ demonstrate the outer and inner radius of aperture, respectively [18]. The purity of Bessel beam can be determined by the size of the inner and outer radius of the annular mask, wavelength and the numerical aperture of the Fourier lens.

An Airy beam can be generated in one or in two-dimensional shapes. Theoretically it can be proved that the Fourier transform of the optical beam with a Gaussian amplitude and cubic phase produces an Airy beam. If the cubic phase just has k-vector in one direction 
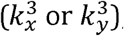
, a 1D Airy beam is achievable. In addition, with superposition of two cubic phase maps in two directions 
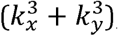
, 2D Airy beam can be obtained. Therefore, the transmittance function of an element for producing an Airy beam can be written as,

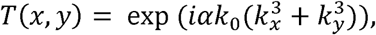

where the dimensionless parameter α (cubic phase coefficient) can defined the property of the Airy beam. In the other words, invariant propagation length of the Airy can be controlled by changing the α [19].

### Damman Grating

An array generator is an optical element that splits the incoming beam to the equal intensity beams. Many different methods were introduced for this purpose. One of the most powerful method to generate an array beam is using a Damman grating. A Damman grating is a binary phase grating that works like as a beam splitter in one or two-dimension. If we assume that all the optical elements have a transmission function, hence, the transmission function of a Damman grating has two values −1 and +1 corresponding to the phase values 0 and pi. The Damman grating is described with transitions points *x*_1_, *x*_2_, …*x_n_* that the value of the transitions points related to the number of the desired intensity spots. The sufficient number of the transitions per period of the grating is critical to creating a specified number of the beams. In general, for generating N beams, one period of the grating needs a 
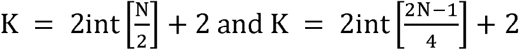
 (K is the transition number), for odd and even number spot, respectively. For this approach, we used the transition points that are calculated with C. Zhou. et al. [13] (Suppl. Table 1). Damman grating makes an incoherent array of the beams and it does not control the relationship between the beams, therefore, for making an array of the beams without superposition the period of the grating should be large enough. It should be noted that the efficiency of the Damman grating in odd numbers is higher than the even numbers. Hence, the odd numbers cover a wide range of applications.

### Optimal Grating

The optimal grating is another method for generating an array of the beams with uniform distribution [15]. The phase function of the optimal grating can be described as:

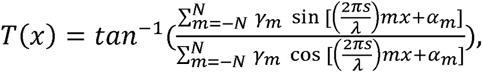

Where the m is the diffraction order, x shows the transverse coordinate, the γand α are relative phase and intensity parameters associated with different diffraction orders which angular separation between them is associated to [15,16]. This method is limited to the odd number of the array intensity. It can support the 3, 5, 7, 9 and 11 fan of the intensity. The space between the beams can be controlled with the period of the grating. In SSPIM toolbox, phase and intensity parameters that are proofed in ref [15] are used, (Suppl. Table 2).

### Lattice beam

Optical lattice is a coherent array beam, which can be generated in one, two, and three dimensions. The SPIM method requires a 1D-like lattice beam as an illumination beam [8]. The 1D optical lattice can be employed for two modes of the light sheet, dithering (scanning) or structured illumination (SIM) mode. This beam can be formed using Bravais pattern. The Bravais pattern is a known 2D lattice defined by the discrete translation operation that can be written as:

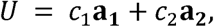

where *c*_1_ and *c*_2_ can get any integer value and **a**_1_ and **a**_2_ are the primitive vectors. The discrete point with this translation operation can make a periodic pattern. The shape of the periodic pattern is related to the size and orientation of the primitive vectors. In general, a Bravias 2D lattice contains five periodic structures including oblique, rectangular, cantered rectangular, square and hexagonal. In order to generating an optical lattice, the first step is to select the parameters of the lattice. These parameters are included the size and the spatial separation of the spots of the Lattice. In the second step, an arbitrary bounding function can confine the lattice in one dimension. Then the spatial frequency and behaviour of the bound lattice can be adjusted with an annular mask in Fourier domain. In the SPIM setup, if the SLM is conjugated to the front focal plane of the objective lens, the SLM pattern is binarized bound lattice. Otherwise, the SLM pattern is the combination of the annular mask and Fourier transform of the bound lattice. The square and hexagonal patterns commonly used as lattice patterns. The lattice beams can be used in different modes, dithering or SIM mode. These two modes can be used for two different reasons such as light confinement and axial resolution. The optimum light confinement can be achieved with control the distances between the cores of the coherent Bessel beams (spatial separation of the spots of the Lattice). On the other hand, the tail of the Bessel beams in some regions produce destructive interference. Hence, the light will be confined in the center of the beam and the out-of-focused will be reduced.

### Tiling method

High-resolution imaging over a large FOV is possible with tiling method [10-11], as well. In this method, the illumination beam with minimum waist (high axial resolution) and small FOV can be swiped in the propagation direction of the illumination light which can make a virtual thin light sheet over a large FOV. For this reason, we need a fast optical element like as tunable lens to tile (swipe) the focused light at the different positions. In the other hand, the tunable lens is able to change the curvature of the incident phase map. Hence, the initial illumination beam such as Gaussian, Bessel or Lattice should be merged with two or more lens phase maps. As mentioned above, the phase map of a lens can be written as:

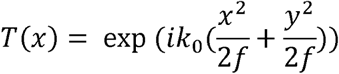

In order to change the position or tiling of a focused light, the focal length of the lens needs to be changed. If we assume, we need three steps tiling to cover the specific FOV, we should select three different focal lengths (*f*_1_,*f*_2_,*f*_3_). Then, we will have three different phase maps (*T*_1_,*T*_2_,*T*_3_) which can be merged with the initial phase map. In total, we will have three phase maps or phase patterns, which can be loaded into the SLM. Each of these beams covers one-third of the field of view. Hence, in total we will have three images from one focus plane which can be merged together.

## Acknowledgement

We thank to Kees Weijer (Dundee, U.K.) and Thomas Baumert (Kassel, Germany) for valuable comments on the manuscript.

## Author Contributions

M. Aakhte and H. A. J. Müller conceived the idea designed toolbox and experiment. M. Aakhte, E. A. Akhlaghi and H. A. J. Müller developed the SSPIM toolbox; M. Aakhte performed experiments and analyzed data; M. Aakhte, E. A. Akhlaghi and H.-Arno J. Müller wrote the manuscript, all authors discussed results and critically revised the manuscript.

## Availability of materials and data

The SSPIM toolbox and the associated documentation are available at https://github.com/aakhtemostafa/SSPIM.

